# Metagenomic analysis of age-dependent microbial dynamics in dual-media rapid sand filters treating groundwater

**DOI:** 10.1101/2024.12.25.630300

**Authors:** Alje S. Boersma, Signe Haukelidsaeter, Francesca Naletto, Caroline P. Slomp, Paul W.J.J. van der Wielen, Maartje A.H.J. van Kessel, Sebastian Lücker

## Abstract

Biofiltration using rapid sand filters is an often used and highly effective method to produce drinking water from anoxic groundwater, yet the microbial processes underlying contaminant removal remain poorly understood. This study used metagenomic sequencing to analyze the microbial communities of dual-media filters of varying ages, yielding 34 high-quality and 177 medium-quality metagenome-assembled genomes (MAGs), including 102 MAGs harboring genes involved in methane, iron or manganese oxidation, and nitrification. Methanotrophic bacteria from the *Methylomonadaceae* family were abundant in both the youngest and oldest anthracite layers. The dominant methanotroph shifted from a MAG possessing both soluble and particulate methane monooxygenases in the two-month-old filters to MAGs only containing particulate methane monooxygenases in older filters, possibly due to changes in copper availability. Iron oxidation was restricted to the anthracite layer and linked predominantly to *Gallionella*, which encoded cluster 1 outer membrane cytochromes Cyc2. *Gallionella* MAGs also became more abundant with filter age, indicating increasing relevance of biological iron oxidation in mature filters. The nitrifying community was dominated by *Nitrosomonas* and Candidatus *Nitrotoga* in the two-month-old filter, with the latter being replaced by *Nitrospira* as the most abundant NOB in older filters. Canonical *Nitrospira* far exceeded comammox *Nitrospira* in abundance, except in the oldest sand layers. These results challenge the previously posed assumption that *Nitrospira* dominance in rapid sand filters is primarily due to comammox activity. Furthermore, it indicates that long operational times are required for comammox *Nitrospira* to achieve significant abundance. Intriguingly, *Nitrospira* MAGs contained multi-copper oxidase-type putative manganese oxidases and cluster 3 *cyc2* genes. Phylogenetic analysis could not conclusively link the *Nitrospira* Cyc2 to iron or manganese oxidation, warranting further study of their role in the removal of these compounds. Overall, these findings reveal dynamic, age-dependent changes in microbial communities and their functions within rapid sand filters, offering new targets for physiological studies of bacteria involved in contaminant removal from anoxic groundwater during drinking water production.

**HIGHLIGHTS:**

- Methanotrophic community shifts between young and mature rapid sand filters.
- Cluster 1 Cyc2-encoding *Gallionella* mediate biological iron oxidation.
- *Candidatus* Nitrotoga and *Nitrosomonas* dominate nitrification in young filters.
- Comammox *Nitrospira* become abundant after prolonged operational periods.
- Canonical *Nitrospira*, not comammox, constitute high *Nitrospira* abundances.
- Abundant cluster 3 Cyc2-containing *Nitrospira* might catalyze metal oxidation.

## 1. INTRODUCTION

In the Netherlands, approximately 60 % of drinking water is sourced from groundwater (Versteegh and Dik, 2012). Groundwater often contains dissolved methane, iron, ammonium, and manganese, all of which must be removed during water treatment. If iron and manganese are not adequately removed, they can cause water discoloration, fouling of distribution systems and household fixtures, give a metallic taste, and, in the case of manganese, pose potential health risks (Vries et al., 2017; WHO, 2017). The removal of methane, ammonium and biodegradable dissolved organic carbon (BDOC) is also crucial, as their presence can promote bacterial regrowth within the distribution network, including potentially pathogenic bacteria (Poghosyan et al., 2017; Volk and Lechevallier, 2002). Through a combination of aeration and rapid sand filtration, these contaminants are effectively removed by abiotic and biological processes, allowing for the safe distribution of drinking water in the Netherlands without the need for residual disinfection, which mitigates associated risks such as toxic trihalomethane production (Smeets et al., 2009).

While most methane is effectively removed during aeration, residual methane is oxidized by methane-oxidizing bacteria, which are ubiquitously present in rapid sand filters (Albers et al., 2015; Boersma et al., 2024; Gülay et al., 2016; Haukelidsaeter et al., 2024; Palomo et al., 2016). Methane oxidation to methanol can be performed by soluble (sMMO) or particulate methane monooxygenase (pMMO; Stein et al., 2012), the latter of which is a member of the copper membrane monooxygenase (CuMMO) enzyme family. Additionally, a sequence-divergent member of the CuMMO enzyme family (pXMO) also might be involved in methane oxidation (Tavormina et al., 2011).

Ammonium is removed exclusively through biological processes. Ammonia-oxidizing bacteria (AOB) such as *Nitrosomonas* and nitrite-oxidizing bacteria (NOB) like *Candidatus* Nitrotoga and *Nitrospira* work in tandem to fully oxidize ammonia to nitrate via nitrite. While ammonia-oxidizing archaea (AOA) are generally rare in rapid sand filters, they are sometimes found in high abundance (van der Wielen et al., 2009). Additionally, they may be present in higher abundance in household filters, where they can contribute to ammonium removal (Van Le et al., 2022). Ammonia is first oxidized to hydroxylamine by ammonia monooxygenase (AMO), another member of the CuMMO family (Musiani et al., 2020). Hydroxylamine is then oxidized to nitric oxide by hydroxylamine oxidoreductase (HAO), which is then converted to nitrite by an unknown enzyme (Caranto and Lancaster, 2017). The produced nitrite is oxidized by NOB to nitrate using the nitrite oxidoreductase (NXR; Chicano et al., 2021).

In rapid sand filters, *Nitrospira* has often been identified as the dominant nitrifier, frequently surpassing the expected ratios of AOB to NOB, assuming a solely nitrite-oxidizing metabolism for *Nitrospira* (Winkler et al., 2012). After the discovery of complete ammonia-oxidizing (comammox) *Nitrospira* (Daims et al., 2015; van Kessel et al., 2015), metagenomic analyses revealed that comammox *Nitrospira* are present in rapid sand filters (Palomo et al., 2016), where they were in some cases found to be the dominant nitrifier (Fowler et al., 2018; Hu et al., 2020; Poghosyan et al., 2020). As a result, it has been suggested that the historical dominance of *Nitrospira* in rapid sand filters may, in part, be attributed to their ability to perform complete ammonia oxidation (Albers et al., 2018; Haukelidsaeter et al., 2023; Palomo et al., 2016; Pinto et al., 2016). However, these studies often did not distinguish between comammox and nitrite-oxidizing *Nitrospira* (hereafter referred to as canonical *Nitrospira*). Consequently, further research is necessary to clarify under what conditions comammox or canonical *Nitrospira* may dominate in rapid sand filters.

Iron and manganese can be removed through both chemical and biological processes. For iron, chemical oxidation in rapid sand filters can occur via both homogeneous and heterogeneous mechanisms (Vries et al., 2017). Biological iron oxidation is performed by iron-oxidizing bacteria, of which *Gallionella* and sometimes *Sideroxydans* are the most commonly found in rapid sand filters treating groundwater (Albers et al., 2015; Müller et al., 2024, Haukelidsaeter et al., 2024). Under neutral aerobic conditions, as found in rapid sand filters, the initial oxidation of iron(II) to iron(III) could be catalyzed by outer membrane *c*-type cytochromes Cyc2 or MtoA (Barco et al., 2015; Liu et al., 2012).

Chemical manganese removal only occurs via heterogeneous oxidation in rapid sand filters (Vries et al., 2017). It was assumed that biological manganese oxidation played a significant role only in young filters, while heterogeneous oxidation became the dominant process in older filters (Bruins et al., 2015). However, recent work suggests biological oxidation remains the primary removal mechanism in rapid sand filters (Haukelidsaeter et al., 2024). Yet, little is known about the specific bacteria responsible for manganese oxidation in rapid sand filters. Potential candidates include bacteria from different genera. For instance, *Pseudomonas* spp. have been linked to manganese oxidation through both culture-dependent and independent methods (Bruins et al., 2017; Hu et al., 2020; Marcus et al., 2017). Other groups, such as *Burkholderiales* and *Rhizobiales*, might be involved since metagenome-assembled genomes (MAGs) of representatives of these genera encode for proteins putatively involved in manganese oxidation (Palomo et al., 2016). Additionally, the rhizobial genera *Pedomicrobium* and *Hyphomicrobium* have been hypothesized to contribute to manganese removal based on their high abundance in rapid sand filters and their capacity to oxidize manganese in drinking water distribution systems and lake sediments (Larsen et al., 1999; Albers et al., 2015; Palermo & Dittrich, 2016). Furthermore, *Leptothrix* might be potentially involved in both manganese and iron oxidation in rapid sand filters (Boogerd and de Vrind, 1987; Fleming et al., 2018). Bacterial manganese oxidation relies on several enzymes, most belonging to the multicopper oxidase (MCO) family. Among these, the manganese-oxidizing multicopper oxidases McoA and MnxG have been identified in *Pseudomonas putida* (Geszvain et al., 2013), with MnxG also found to oxidize manganese in *Bacillus* (Dick et al., 2008). In *Bacillus*, manganese oxidation can additionally be supported by the spore coat protein A (CotA; Su et al., 2013). MoxA, identified in *Pedomicrobium*, and MofA, identified in *Leptothrix discophora* are additional MCOs essential for manganese oxidation activity in these bacteria (Ridge et al., 2007; Corstjens et al., 1996). Besides MCOs, other enzyme families contribute to manganese oxidation. Peroxidase-cyclooxygenases, also referred to as animal heme peroxidases, can oxidize manganese directly, as seen in *Erythrobacter* and *Aurantimonas* (Anderson et al., 2009; Medina et al., 2018), or indirectly via superoxide production, as observed in *Roseobacter* (Andeer et al., 2015).

The physiological role of manganese oxidation by MCOs and perixodase-cyclooxygenases is, however, still unknown. Firstly, despite manganese oxidation being thermodynamically favorable, its use for energy conservation has not been shown (Geszvain et al., 2012). An alternative role may involve protecting cells from reactive oxygen species or free radicals, which manganese oxides can neutralize (Daly, 2009). Additionally, manganese oxides can oxidize refractory organic compounds, potentially enabling bacteria to utilize them as energy and carbon sources (Tebo et al., 2004). This aligns with the observation that MCOs are evolutionarily related to laccases involved in lignin degradation (de Gonzalo et al., 2016) and can even exhibit laccase activity themselves (Su et al., 2013; Ridge et al., 2007). Recently, the first instance of energy conservation through manganese oxidation was demonstrated in *Candidatus* Manganitrophus noduliformans. This chemolithoautotrophic bacterium, a member of the phylum *Nitrospirota* and a distant relative of nitrifying *Nitrospira*, is hypothesized to use an iron-oxidizing outer membrane *c*-type cytochrome Cyc2 homolog as the initial oxidant and electron carrier for manganese oxidation (Yu and Leadbetter, 2020) Notably, while these enzymes exhibit manganese-oxidizing activity, none of the associated organisms have been isolated from rapid sand filters and, consequently, the study of manganese oxidation in these systems remains limited to the investigation of homologous genes.

Here, we investigated the microbial community of rapid sand filters of varying ages using metagenomics. We aim to enhance the understanding of the role microorganisms play in contaminant removal, and to identify bacteria that were previously not associated with this process. Since biological oxidation plays a key role in the removal of methane, iron, ammonium and manganese in rapid sand filters, the responsible organisms must be identified to create a comprehensive understanding of rapid sand filtration. Employing both gene and genome (MAG)-centric approaches, we studied the role of bacteria in methane, iron, ammonium, and manganese removal, focusing mostly on the latter two.

## 2. MATERIALS AND METHODS

### 2.1 Drinking water treatment plant

Samples were collected from the Vitens N.V. drinking water treatment plant in Sint Jansklooster, The Netherlands, where groundwater is treated in an 11-step process, as detailed by Haukelidsaeter et al. (2023). The chemical composition of the raw water at this site varies due to the alternating extraction of groundwater from 18 wells. Typically, the methane concentration ranges from 195 to 211 µM (Boersma et al., 2024), iron from 125 to 180 µM, manganese from 6 to 8 µM, and ammonium from 70 to 200 µM. The water temperature generally remains between 11 and 12 °C.

This study focuses on the primary rapid sand filtration step, where most iron, ammonium, and manganese are removed. After initial plate aeration to strip methane, raw water is distributed across 12 parallel dual-media filters, each with a surface area of 25 m^2^. These filters consist of a 0.9 m layer of anthracite (particle size: 1.4–2.5 mm) above a 1.6 m layer of sand (particle size: 0.8–1.2 mm), with porosities of approximately 50 % and 42 %, respectively. The average supernatant depth above the filters is approximately 0.1 m.

### 2.2 Sample collection

We investigated four filters of varying ages. Filter 1 was sampled in March 2021 when the anthracite and sand layers were two and eleven years old, respectively. Filter 2 was sampled in April 2021 when the anthracite and sand layers both were seven months old. Filter 3 was sampled in April and October 2021, when both the anthracite and sand layers were two and seven months old, respectively. Filter 4 was sampled in April 2021, when the anthracite and sand layers were one and eleven years old, respectively.

Samples were taken as described in detail by Haukelidsaeter et al. (2023). Because previous research showed that the anthracite and sand layers contained homogeneous microbial communities as a result of frequent backwashing (Boersma et al., 2024), the 0-4 cm and 4-10 cm depth samples were pooled to create one sample for the anthracite layer, and the 100-150 cm and 150-200 cm samples were pooled to create one sample for the sand layer. The samples analyzed here are thus representative of the entire filter medium layer.

### 2.3 DNA isolation and sequencing

DNA was isolated using 0.5 g (wet weight) of the pooled anthracite or sand samples using the DNeasy Powersoil Kit (QIAGEN, Hilden, Germany). Cell lysis was performed by bead beating at 50 Hz for 1 min using a TissueLyser LT (QIAGEN, Hilden, Germany). Sequencing was performed by Macrogen Inc. (Seoul, South Korea) using the Illumina Novaseq 6000 platform and the S4 flow cell, resulting in 2×150 bp paired-end reads. Paired-end libraries were constructed using the Nextera XT DNA Library Preparation Kit (Illumina, San Diego, USA) with the Nextera XT DNA Library Prep Kit Reference Guide (15031942 v03).

### 2.6 Metagenome Assembly

Metagenomic paired-end reads were quality-filtered, decontaminated, and adapter-trimmed to a minimum length of 75 bp using BBDuk (BBTools v37.76, https://jgi.doe.gov/data-and-tools/software-tools/bbtools/). Trimmed reads from the anthracite and sand layer of each filter and sampling time were co-assembled with metaSPAdes v3.15.4 (Nurk et al., 2017). Assembly was conducted with k-mer sizes of 21, 33, 55, 77, and 99, retaining only contigs ≥ 1000 bp for further analysis.

### 2.7 Metagenome Binning

Reads from each sample were mapped to the assemblies using BBmap (BBTools v37.76) to generate differential coverage data, which was converted to BAM format via SAMtools v1.8 (Li et al., 2009). Binning was performed using BinSanity v0.2.6.3 (Graham et al., 2017), CONCOCT v0.4.1 (Alneberg et al., 2013), MaxBin2 v2.2.7 (Wu et al., 2016), and MetaBat2 v2.12.1 (Kang et al., 2019), with consensus binning by DAS Tool v1.1.1 (Sieber et al., 2018). Bins were dereplicated with dRep (Olm et al., 2017) at 99% ANI, then refined manually in Anvi’o v7.1 (Eren et al., 2021). Completeness and contamination of bins were assessed with CheckM2 v0.1.3 (Chklovski et al., 2023) and classified according to MIMAG standards (Bowers et al., 2017). Taxonomy was assigned using GTDBtk v2.1.1 (Chaumeil et al., 2022), and relative abundance was calculated with CoverM v0.4.0 (https://github.com/wwood/CoverM).

### 2.8 Gene Annotation

Gene annotation was conducted using DRAM v1.2.3 (Shaffer et al., 2020), with iron-cycling genes predicted by FeGenie (Garber et al., 2020). To enhance nitrogen-cycling gene identification, DRAM-predicted open reading frames (ORFs) generated by Prodigal (Hyatt et al., 2010) were mined for nitrogen cycle genes using Metascan HMMs (Cremers et al., 2022) to identify additional *nxrA* genes. Gene coverage was computed with CoverM v0.4.0, using RPKM normalization for gene size and sequencing depth.

### 2.9 Phylogenetic Analysis

The phylogeny of *Nitrospira* MAGs identified in this study and a reference database of 265 genomes was inferred with the UBCG pipeline by extracting, aligning, and concatenating 92 single-copy core genes (Na et al., 2018). A maximum-likelihood tree was generated from this alignment with IQ-TREE v2.1.4 (Minh et al., 2020) using the GTR+F+I+G4 model and 1000 ultrafast bootstraps. 59 CuMMO sequences were aligned with 490 reference sequences using MUSCLE v3.8.1551 (Edgar, 2004), positions with more than 50 % gaps were removed using ClipKIT v1.3.0 (Steenwyk et al., 2020), and a maximum-likelihood tree was built using IQ-TREE v2.1.4 (Minh et al., 2020) with the LG+F+I+G4 model and 1000 ultrafast bootstraps. For determining the phylogeny of NxrA and NarG, 202 sequences and a reference set of 423 sequences (Poghosyan et al., 2020) were aligned with ARB v5.5 (Ludwig et al., 2004). A maximum-likelihood tree was generated in IQ-TREE v2.1.4 (Minh et al., 2020) using the Q.pfam+I+G4 model and 1000 ultrafast bootstraps. Phylogeny of Cyc2 was investigated by aligning 32 sequences to a reference alignment of 1593 sequences (Keffer et al., 2021), which also included the Cyc2 from *Candidatus* Manganitrophus noduliformans (Yu and Leadbetter, 2020). This alignment was performed using MUSCLE v3.8.1551 (Edgar, 2004), positions with more than 30 % gaps were removed with ClipKIT v1.3.0 (Steenwyk et al., 2020), and a maximum-likelihood tree was calculated with IQ-TREE v2.1.4 (Minh et al., 2020) using the VT+F+I+G4 model and 1000 ultrafast bootstraps. All trees were visualized with iTOL (Letunic and Bork, 2021).

## 3. RESULTS & DISCUSSION

Metagenomic sequencing yielded 73-86 million paired-end raw reads, with 67-80 million reads left after trimming. Of the trimmed reads, 61-82 % were assembled into contigs, of which 68-79 % were binned into MAGs (Table 1). This resulted in 34 high (≥90 % completeness, <5 % contamination, a complete small subunit rRNA operon and ≥18 tRNAs) and 177 medium-quality (≥70 % completeness and <10 % contamination; Table S1). Of these 211 MAGs, 102 MAGs stemmed from bacteria possibly involved in contaminant removal as they contained genes possibly involved in methane, iron or manganese oxidation, or nitrification (Figure 1). Removal profiles of iron, ammonium, and manganese and the effect of filter age were discussed in detail by Haukelidsaeter et al. (2023). In short, two months after startup, Filter 3 failed to remove ammonium and manganese due to a lack of biofilm and mineral coating on the sand grains. Filters of intermediate age (Filter 2 at seven months and Filter 3 at eight months) removed ammonium and manganese optimally. The old Filters 1 (two and eleven-year-old anthracite and sand, respectively) and 4 (one and eleven-year-old anthracite and sand, respectively) removed all ammonium but failed to completely remove manganese, likely due to preferential flow of water through the filter. Lastly, since the anthracite and sand layers are homogenous as a result of backwashing (Boersma et al., 2024), the metagenomes represent the entire filter medium layer.

**Figure 1.**
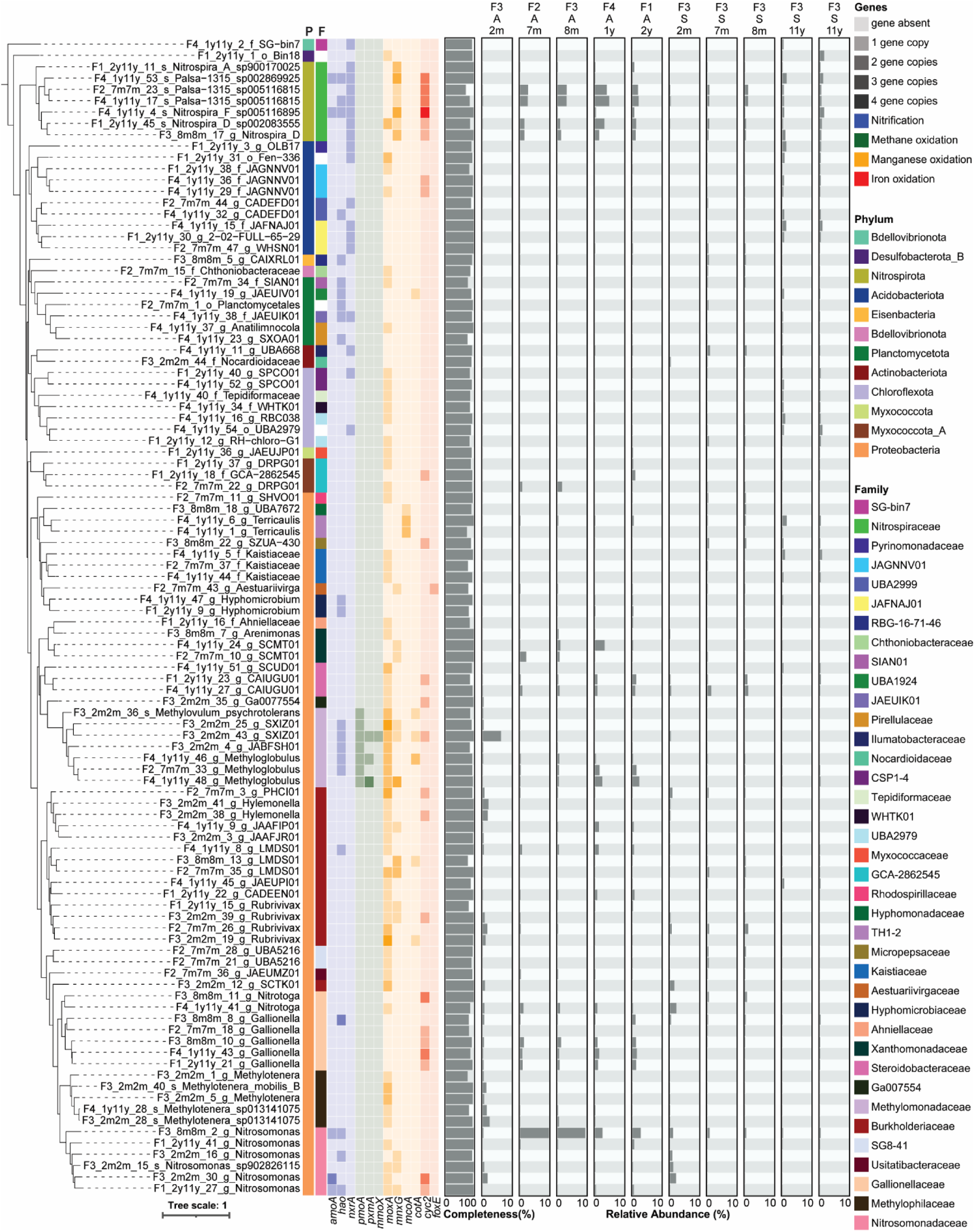
Overview of phylogeny, taxonomy, genetic potential, completeness and distribution of 102 MAGs containing genes involved in methane, iron and manganese oxidation, and nitrification. The MAGs were selected out of 211 total MAGs based on the presence of genes involved in methane, iron or manganese oxidation, and nitrification. The MAGs are ordered by phylogeny, as determined by a maximum likelihood tree made with IQ-tree, using the GTR+F+I+G4 model, based on an alignment made by UBCG, and rooted using three archaeal sequences as outgroup. Taxonomy of the Phylum (P) and Family (F) level is shown in colored bars next to the phylogenetic tree. Copy numbers of genes involved in nitrification, methane, iron, and manganese removal are shown as a heatmap. Bar plots show MAG completeness as calculated by checkm2 and relative abundance across all samples calculated by CoverM. The samples are separated based on filter medium and ordered in ascending age. A = anthracite, S = sand, m = month, y = year.

**Table 1.** Statistics of metagenomics analysis per sample. Filter name is followed by age of filter medium and filter medium type, y = year, m = month, A = anthracite, S = sand.

| Statistic | F1 A 1y | F1 S 11y | F2 A 7m | F2 S 7m | F3 A 2m | F3 S 2m | F3 A 8m | F3 S 8m | F4 A 2y | F4 S 11y |
| --- | --- | --- | --- | --- | --- | --- | --- | --- | --- | --- |
| Total reads | 78,139,182 | 78,235,822 | 83,369,150 | 78,318,414 | 80,910,446 | 79,076,628 | 73,252,214 | 77,679,306 | 82,290,472 | 85,722,298 |
| Trimmed reads | 71,143,600 | 71,315,366 | 76,208,770 | 69,246,990 | 71,670,968 | 74,728,842 | 67,038,868 | 69,276,378 | 73,569,418 | 79,709,576 |
| Assembled trimmed reads (%)<sup>a</sup> | 74.62 | 63.90 | 81.14 | 66.88 | 81.61 | 81.52 | 77.81 | 60.83 | 81.20 | 71.73 |
| Assembled trimmed reads in MAGs (%)<sup>b</sup> | 73.48 | 70.93 | 74.31 | 72.46 | 79.00 | 76.01 | 72.68 | 68.40 | 77.10 | 74.60 |
| Number of contigs | 273,028 |  | 232,629 |  | 137,386 |  | 197,060 |  | 277,155 |  |
| Longest contig | 484,855 |  | 955,811 |  | 619,238 |  | 652,304 |  | 1,139,809 |  |
| N50 | 3526 |  | 3844 |  | 4518 |  | 3377 |  | 3895 |  |

<sup>a</sup>Determined by BBTools BBMap v37.76 (https://jgi.doe.gov/data-and-tools/software-tools/bbtools/)
<sup>b</sup>Determined by CoverM v0.4.0 (https://github.com/wwood/CoverM)

### 3.1 Methane-oxidizing bacteria are abundant but composition changes as filters age

All MAGs harboring genes associated with methane oxidation belong to the *Methylomonadaceae* family (Figure 1). These MAGs are abundant in the anthracite layer across all filter ages, except at 7 and 8 months. However, their distribution shifts as filters age. At two months, the most abundant methanotrophic MAG in the anthracite layer is F3_2m2m_43 (8.9 %), which contains not only the genes encoding the membrane-bound pMMO (*pmoA*) and pXMO (*pxmA*) but also the sMMO (*mmoX*). Over time, F3_2m2m_43 becomes nearly undetectable (0.005 %), while F4_1y11y_48 and F2_7m7m_33 emerge as the most abundant methanotrophic MAGs (3.3–3.6 % and 2.0–2.2 %, respectively). Both of these MAGs contain one *pmoA* gene, and F2_7m7m_33 additionally carries two copies of *pxmA*.

The dominance of F3_2m2m_43 in the young filter (two months) raises the question of whether the presence of a sMMO could explain its early abundance. The sMMO is typically expressed under copper-limited conditions, as copper serves as a cofactor for pMMO (Banerjee et al., 2024). Although copper concentrations were not measured in this study, there is no obvious reason to anticipate more copper limitation at two months compared to later stages based solely on dissolved copper levels. Conversely, ammonium removal was incomplete at this stage (Haukelidsaeter et al., 2023), and since copper is also a cofactor for AMO, it is plausible that copper availability was higher in young filters. However, this does not account for the potential accumulation of copper on sand grains. Extracellular polymeric substances (EPS) in biofilms and mineral coatings can bind copper (Tebo et al., 2004; Zhang et al., 2024), suggesting that copper availability might increase with filter age as both biofilms and mineral coatings become more abundant (Haukelidsaeter et al., 2023). It has also been proposed that sMMO-containing methanotrophs may be favored in environments with high iron concentrations due to the diiron cluster at the active site of sMMO (Poghosyan et al., 2020). This could explain the early dominance of a sMMO-encoding MAG in young filters, where the lack of biofilm and mineral-associated copper would favor sMMO under relatively copper-limited but iron-replete conditions.

The gene-centric analysis, which includes genes not contained in any retrieved MAG, reinforces the presence of *mmoX* exclusively in the two-month-old anthracite layer (Figure 2). However, *pmoA* was consistently the most abundant methane-oxidizing gene at all filter ages followed by *pxmA*, indicating that copper-dependent methane oxidation was the dominant removal process.

**Figure 2.**
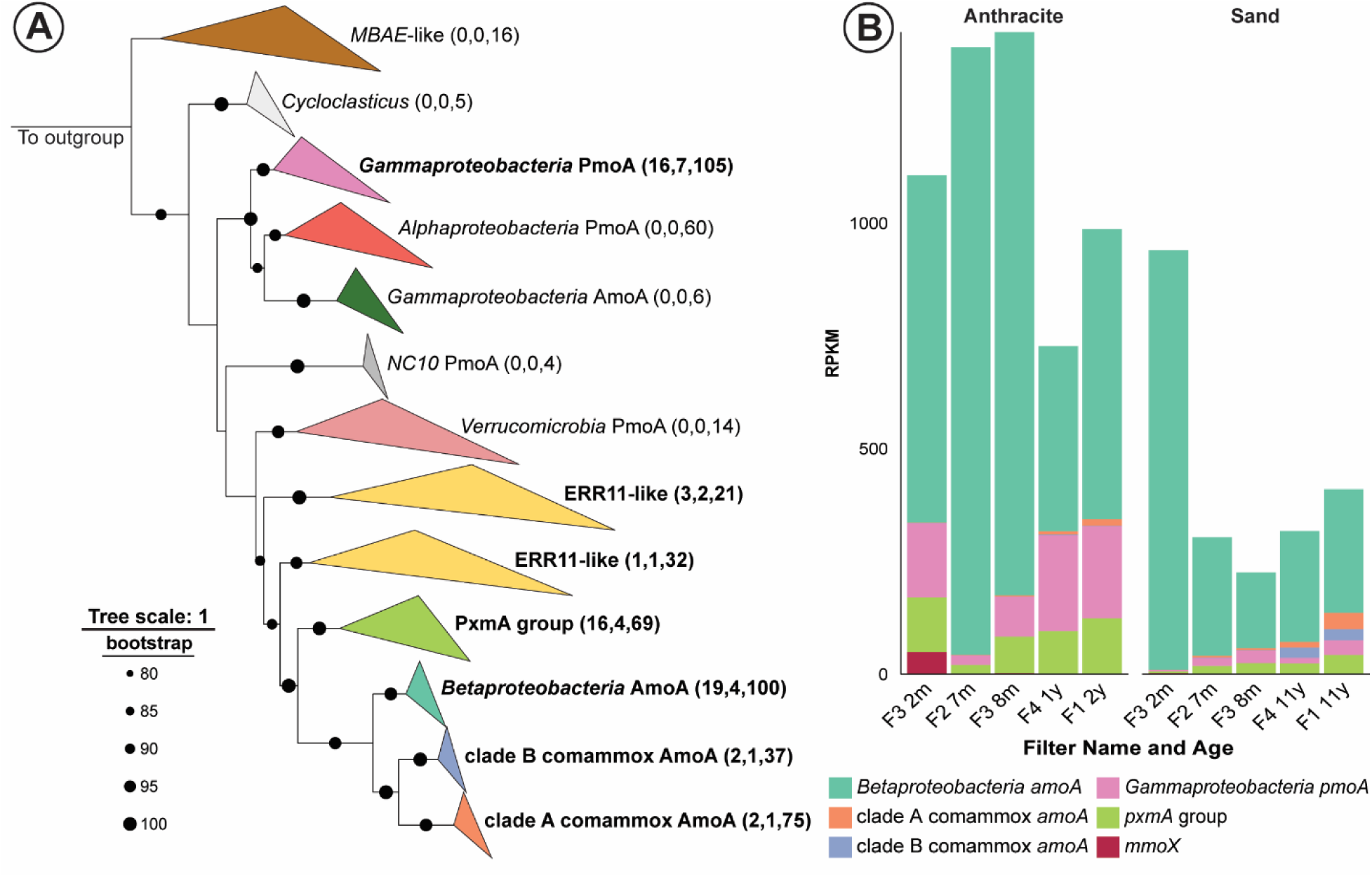
Diversity and abundance of proteins and genes involved in methane and ammonia oxidation. (A) Phylogenetic tree illustrating the diversity of copper-containing membrane-bound monooxygenase-encoding (Cu MMO) proteins. Clades highlighted in bold represent sequences derived from the rapid sand filter metagenomes analyzed in this study. Numbers in brackets indicate the count of sequences from the rapid sand filter metagenomes, the number of these sequences identified within MAGs, and the total number of sequences in the clade, respectively. The tree is rooted with three *Thaumarchaeota* AmoA sequences serving as the outgroup. (B) Relative abundances of the betaproteobacterial *amoA*, clade A and clade B comammox *Nitrospira amoA*, and *gammaproteobacterial pmoA* and *pxmA* genes across all filters, expressed as RPKM (reads per kilobase of gene per million mapped reads). Although not part of the Cu-MMO family, the *mmoX* gene is included as the gene encoding soluble methane monooxygenase (sMMO). The samples are separated based on filter medium and ordered in ascending age. m = month and y = year. A vector image of the full tree can be found in the supplementary materials (https://doi.org/10.5281/zenodo.14554862)

### 3.2 Biological iron oxidation is performed exclusively by cluster 1 Cyc2-containing iron oxidizers

The gene *cyc2* was the most abundant iron-oxidizing gene identified in the MAGs generated in this study (Figure 1). Among these *cyc2-*contaning MAGs, the only ones with taxonomy known to be involved in iron oxidation were members of the genus *Gallionella.* In addition, one of the MAGs (F3_8m8m_8) did not contain *cyc2*. However, as *cyc2* is found in nearly all characterized neutrophilic iron-oxidizing bacteria, including *Gallionella* (Hoover et al., 2023), and F3_8m8m_8 has only 82.11 % completeness, it is plausible that *cyc2* is absent due to incomplete assembly.

Contrastingly, the decaheme *c*-type cytochrome MtoA was not identified in the sand filter metagenomes, but one MAG of the genus *Aestuariivirga* (F2_7m7m_43) encoded FoxE, an enzyme responsible for iron(II) oxidation in photosynthetic iron-oxidizing bacteria (Pereira et al., 2017). The dark conditions in the sand filters studied here do not support the presence of photosynthetic organisms. Furthermore, characterized members of the genus *Aestuariivirga* are known to be aerobic heterotrophs capable of degrading various carbon compounds and are not associated with photosynthesis (Lemos et al., 2021). It is therefore likely that the *foxE* gene identified in this MAG is misannotated and serves a different function. Instead, this MAG is likely involved in BDOC degradation, although its contribution appears minimal, as indicated by its low abundance across all samples (0.001–0.2%).

Interestingly, Cyc2 homologs were also identified in MAGs that belong to taxa not traditionally associated with iron oxidation (Figure 1). Phylogenetic analysis reveals that Cyc2 is distributed into three distinct clusters (1, 2, and 3; McAllister et al., 2020). Most Cyc2 sequences found in neutrophilic iron-oxidizing bacteria belong to cluster 1, suggesting a common function in iron oxidation for members of this cluster (McAllister et al., 2020). Additionally, iron-oxidizing activity in cluster 1 Cyc2 was biochemically verified recently for the first time (Keffer et al., 2021). All the *cyc2* genes from *Gallionella* MAGs identified here belonged to cluster 1. Three additional MAGs (F2_7m7m_3, F3_2m2m_39, F3_8m8m_11) assigned to the genera PHCI01*, Rubrivivax*, and *Candidatus* Nitrotoga, respectively, contained cluster 1 Cyc2 sequences. While none of these genera are known to oxidize iron, the presence of cluster 1 Cyc2 in these MAGs warrants further investigation into their potential role in iron oxidation.

Clusters 2 and 3 also each contain one Cyc2 homolog with verified iron-oxidizing activity (Castelle et al., 2008; Jeans et al., 2008). Since all three Cyc2 clusters only contain one functionally verified iron-oxidizing sequence, care must be taken when attributing iron-oxidizing function to various Cyc2 homologs. A good predictor of function is how closely related the protein is to a characterized Cyc2, or if a protein sequence falls within a well-supported clade of known iron-oxidizing bacteria, such as *Gallionella* (Keffer et al., 2021). Especially for cluster 3, its function can be considered uncertain, as the only functionally characterized sequence is phylogenetically distant from other sequences in the clade, indicating that it is not necessarily representative of this cluster (Keffer et al., 2021).

In conclusion, Cyc2 was the predominant iron-oxidizing enzyme detected in the sand filter metagenomes, and cluster 1 Cyc2-containing *Gallionella* were responsible for biological iron oxidation. *Gallionella* MAGs were most abundant in the anthracite layer, where their relative abundance increased from 2.3 % at a filter age of two months old to 6.9 % at two years. As total bacterial abundances as measured by qPCR also increased during this time (Haukelidsaeter et al., 2023), this indicates that biological iron oxidation became more relevant in older filters. *Gallionella* are often found in high abundance in rapid sand filters based on 16S rRNA gene amplicon sequencing (Albers et al., 2018; Corbera-Rubio et al., 2024breehei; Haukelidsaeter et al., 2023; Haukelidsaeter et al., 2024), where they are linked to iron oxidation based on taxonomy. The presence of cluster 1 Cyc2 in the *Gallionella* MAGs identified here provides additional evidence for the involvement of *Gallionella* in biological iron oxidation in rapid sand filters.

### 3.3 Canonical *Nitrospira* largely dominate the nitrifying community

The nitrifying community consisted of *Nitrosomonas, Candidatus* Nitrotoga, and *Nitrospira.* All 7 *Nitrospira* MAGs encoded NxrA required for nitrite oxidation. In addition, *Nitrospira* MAGs F4_1y11y_4 and F4_1y11y_53 contained the *amoA* and *hao* genes and are thus capable of complete ammonia oxidation. To account for any *amoA* genes missed in the binning process, we performed a phylogenomic analysis of all *Nitrospira* MAGs obtained, confirming that F4_1y11y_4 and F41y11y_53 were the only two comammox MAGs, with F4_1y11y_4 belonging to comammox clade A and F4_1y11y_53 belonging to clade B (Figure S1).

At a filter age of two months, the most abundant nitrifier MAGs were from the genera *Nitrosomonas* (3.5% in the anthracite and 7.2 % in the sand) and *Candidatus* Nitrotoga (0.9 % in the anthracite and 3.1 % in the sand). As the anthracite matured, nitrifiers increased in relative abundance, reaching 29 % and 30 % at seven and eight months, respectively (Figure S2). *Nitrosomonas* remained the dominant AOB at 14 % during both months, while canonical *Nitrospira* (13 % and 15 %) surpassed *Candidatus* Nitrotoga (1.6 % and 1.2 %) as the dominant NOB. In the one-year-old anthracite, the relative abundance of canonical *Nitrospira* peaked at 20 % before decreasing to 11 % at two years. At these time points, assuming a strictly nitrite-oxidizing metabolism for *Nitrospira*, the ratios of canonical *Nitrospira* to *Nitrosomonas* (4.8 at one year and 1.9 at two years) greatly exceeded the expected NOB:AOB ratio of 0.5 (Winkler et al., 2012). This ratio is derived from the number of electrons obtained by nitrite and ammonia oxidation, which are two and six, respectively. However, due to the concomitant reduction of O_2_ by AMO, four electrons are unavailable for energy conservation, and the number of electrons generated is thus similar for AOB and NOB (Lancaster et al., 2018). Still, AOB can conserve more energy as these electrons enter the respiratory chain at the level of the quinone pool and not via cytochrome *c* as in NOB. Comammox *Nitrospira* did emerge in the anthracite, increasing from 0.5 % at one year to 1.0 % at two years, but remained less abundant than the other nitrifiers.

Only in the newly started filter at an age of two months, nitrifiers had a higher relative abundance in the sand (11 %) than in the anthracite layer (5 %). At this point, they likely were outcompeted by methanotrophs in the anthracite, causing higher ammonium concentrations to enter the sand layer. In seven and eight-month-old filters, nitrifier abundance in the sand decreased to 6 % and 7 %, respectively, with *Nitrosomonas* (1.0 % and 0.8 %) and canonical *Nitrospira* (3 % and 5 %) as the dominant AOB and NOB. In the eleven-year-old filter, nitrifier relative abundance in the sand increased again, reaching 7 % in Filter 1 and 4 % in Filter 4, driven by a rise in comammox *Nitrospira* (4 % in Filter 1 and 3 % in Filter 4), which became the dominant ammonia oxidizer during this period.

Out of the 19 identified betaproteobacterial *amoA* genes, only four were assigned to MAGs, along with half of the comammox *amoA* genes (Figure 2). Notably, unbinned *amoA* genes accounted for 82–91 % of the total betaproteobacterial *amoA* abundance and approximately 50 % of the total comammox *amoA* abundance (Figure S3), thereby significantly contributing to the overall *amoA* gene pool. To comprehensively analyze the distribution of all *amoA* genes, including unbinned ones, a gene-centric approach was employed. This analysis revealed that betaproteobacterial *amoA* sequences were the most abundant in all samples (Figure 2), representing 97–100 % of total *amoA* abundance across all samples except the oldest sand layers, where their contribution was lower, at 82–88 %. The dominance of betaproteobacterial *amoA* over comammox *Nitrospira amoA* was more pronounced than the comparison of *Nitrosomonas* and comammox *Nitrospira* MAG relative abundances, suggesting that *Nitrosomonas* is an even more abundant AOB than indicated by MAG abundance. It is important to note, however, that most comammox organisms harbor one to two copies of *amoA*, whereas most AOB have two to three copies (Hommes et al., 2001; Kikuchi et al., 2023; Koch et al., 2009). This discrepancy may lead to an overestimation of betaproteobacterial AOB abundance when extrapolated directly from gene abundance.

Dissimilatory nitrate reductases (NAR) and nitrite oxidoreductases are closely related enzymes, and phylogenetic analysis of NxrA/NarG sequences is essential to assign putative functions to these sequences. The sequences identified in the rapid sand filter metagenomes were affiliated with six distinct phylogenetic groups (Figure 3). The first two comprised *Nitrotoga* and *Nitrospira* NxrA and are confidently associated with nitrite oxidation (Kitzinger et al., 2018; Lücker et al., 2010). The third included a diverse group of NarG sequences, likely involved in nitrate reduction (Zhao et al., 2023). Finally, three groups, with uncertain functions, were identified and these clustered closely with confirmed NxrA sequences. These included putative *Nitrotoga*-like NxrA/NarG, putative *Chloroflexota*-like NxrA/NarG (Spieck 2020), and putative *Brocadiaceae*-like NxrA/NarG (Zhao et al., 2023). Notably, the high abundance of *Candidatus* Nitrotoga MAGs in the youngest filter was reflected by the dominance of *Nitrotoga*-like *nxrA* at this stage. As the filters aged, the *Nitrospira nxrA* gene became the most abundant sequence type, corroborating the observed increase in *Nitrospira* abundance with filter maturation in the MAG-centric analysis. Although the *nxrA/narG* gene sequences with uncertain functions were of low abundance, the isolation and physiological characterization of bacteria harboring these genes are necessary to elucidate their potential roles within rapid sand filters.

**Figure 3.**
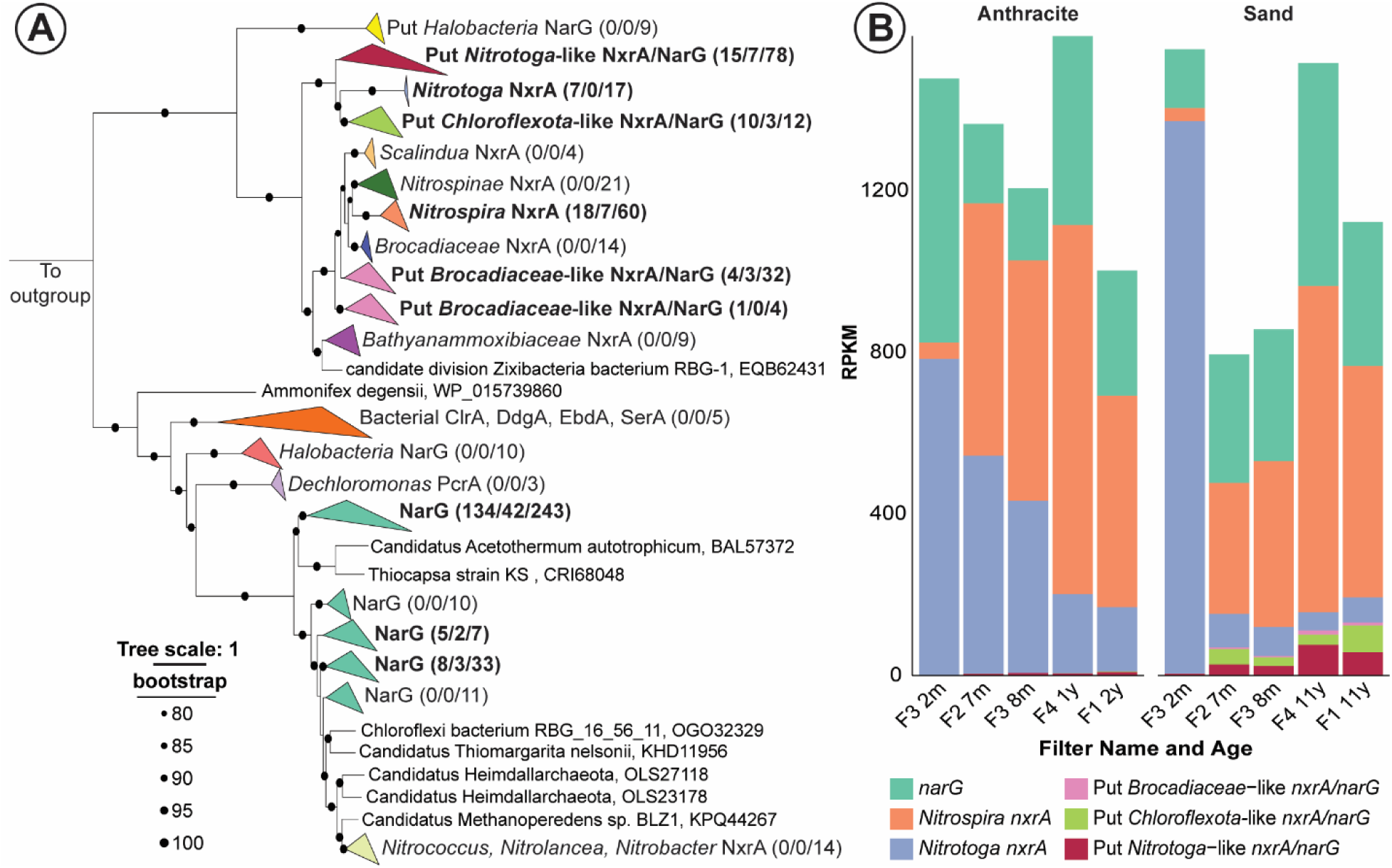
Diversity and abundance of NxrA and NarG. (A) Phylogenetic tree illustrating the diversity of NxrA and NarG proteins. Clades highlighted in bold represent sequences derived from the rapid sand filter metagenomes analyzed in this study. Numbers in brackets indicate the count of sequences from the rapid sand filter metagenomes, the number of these sequences identified within MAGs, and the total number of sequences in the clade, respectively. The tree is rooted with twenty putative PcrA sequences serving as the outgroup. (B) Relative abundances of *narG*, *Nitrospira nxrA*, *Nitrotoga nxrA,* putative *Brocadiaceae-* like *nxrA/narG,* putative *Chloroflexota-*like *nxrA/narG* and putative *Nitrotoga-*like *nxrA/narG* genes across all filters, expressed as RPKM (reads per kilobase of gene per million mapped reads). The samples are separated based on filter medium and ordered in ascending age. Here, m = month and y = year. A vector image of the full tree can be found in the supplementary materials (https://doi.org/10.5281/zenodo.14554862)

Since the discovery of comammox *Nitrospira* (Daims et al., 2015; van Kessel et al., 2015), the high relative abundance of *Nitrospira* compared to *Nitrosomonas* in rapid sand filters has often been attributed to comammox *Nitrospira*, typically without distinguishing between canonical and comammox *Nitrospira* (Albers et al., 2018; Haukelidsaeter et al., 2023; Palomo et al., 2016; Pinto et al., 2016). Our findings challenge this assumption, showing that the abundance of *Nitrospira* in the filters was predominantly due to canonical *Nitrospira*, except in the oldest sand layers. These findings suggest that comammox *Nitrospira* may only become dominant after extended operational periods, but it remains to be studied to what extent our findings can be extrapolated to rapid sand filters of other treatment plants.

Still, our finding is supported by other research that quantified comammox *Nitrospira*, and which found them to be dominant in a filter of a different treatment plant in the Netherlands that was sixteen to eighteen years old at the time of sampling (Poghosyan et al., 2020), but in younger filter (one-year old) at the same location, canonical *Nitrospira* remained the dominant nitrifier (Corbera-Rubio et al., 2024, Breehei). In other research, comammox *Nitrospira* were the dominant AOB in one of three investigated drinking water treatment plants treating groundwater (Hu et al., 2020). However, the ages of the filters were not reported and NOB were not quantified in that study, making it difficult to compare with our research.

### 3.4 A potential role of *Nitrospira* in metal oxidation

As discussed above, canonical *Nitrospira* were the dominant nitrifiers in this study, exhibiting abundances far exceeding the expected NOB:AOB ratio of 0.5, based on a strictly nitrite-oxidizing metabolism. This raises the question of whether *Nitrospira* may participate in processes other than nitrification. Notably, all *Nitrospira* MAGs identified contained one or more copies of the *mnxG* gene, and MAG F1_2y11y_45 additionally harbored a *moxA* gene (Figure 1). These MCOs were widespread across the MAGs in this study, including taxa that are not traditionally linked to manganese oxidation, such as *Nitrosomonas* and *Methylomonadaceae* (Figure 1). However, homology is not a good predictor of function for these genes (Kurdi et al., 2023), and a role for *Nitrospira* or the other taxa in manganese oxidation purely based on the presence of MCO genes can thus not be concluded.

All bacterial MCOs with manganese-oxidizing activity have been identified in heterotrophic bacteria (Kurdi et al., 2023), are closely related to laccases (de Gonzalo et al., 2016), and can have laccase activity themselves (Su et al., 2013; Ridge et al., 2007). As also observed in the MAGs obtained in this study, this might explain the widespread occurrence of these genes in heterotrophic bacteria, where they could be involved in the degradation of phenolic substrates. It might also explain why during isolation attempts of manganese-oxidizing bacteria from rapid sand filters, heterotrophic bacteria such as *Pseudomonas* and *Bacillus* are often identified (Bruins et al., 2017; Marcus et al., 2016), as they can contain MCOs capable of manganese oxidation but could also be involved in heterotrophic metabolisms *in situ*.

So far, members of only one bacterial genus – *Candidatus* Manganitrophus – have been found to perform chemolithoautotrophic manganese oxidation, namely *Ca.* M. noduliformans and *Ca.* M. morganii (Yu and Leadbetter, 2020; Yu et al., 2022). Here, a Cyc2 homolog was expressed and hypothesized to perform the initial oxidation of manganese. Six out of seven *Nitrospira* MAGs identified in this study also encode Cyc2 (Figure 1). Phylogenetic analysis of the Cyc2s of *Ca.* M. noduliformans, in the public database (Keffer et al., 2021), and identified in the MAGs obtained in our study revealed that all Cyc2s of *Nitrospira* as well as *Ca.* M. noduliformans belong to cluster 3 (Figure 4). This phylogenetic assignment was confirmed with FeGenie (Garber et al., 2020), which can distinguish between the different Cyc2 clusters. Interestingly, Cyc2 cluster 3 contains sequences from the manganese-oxidizing *Ca.* M. noduliformans as well as from acidophilic iron-oxidizing bacteria of the genus *Leptospirillum*. However, the *Nitrospira* Cyc2 sequences do not cluster together with neither of these metal oxidizers but, instead, fall within three distinct clades and cluster with other *Nitrospira* sequences. Thus, based on phylogeny, *Nitrospira* Cyc2s could be involved in both iron and manganese oxidation. Such a metal-oxidizing role of *Nitrospira* may explain their consistently high abundances observed in rapid sand filters across various treatment plants (Albers et al., 2018; Corbera-Rubio et al., 2024; Haukelidsaeter et al., 2023; Palomo et al., 2016; Pinto et al., 2016), which cannot be solely sustained by nitrite oxidation given the excess of NOB over AOB. Together with the presence of possible manganese-oxidizing MCOs and Cyc2 homologs in six out of seven *Nitrospira* MAGs retrieved in our study, we hypothesize that *Nitrospira* is involved in metal oxidation in rapid sand filters. Verification of this hypothesis requires the isolation of *Nitrospira* from rapid sand filters and subsequent investigation of their iron and manganese-oxidizing capabilities.

**Figure 4.**
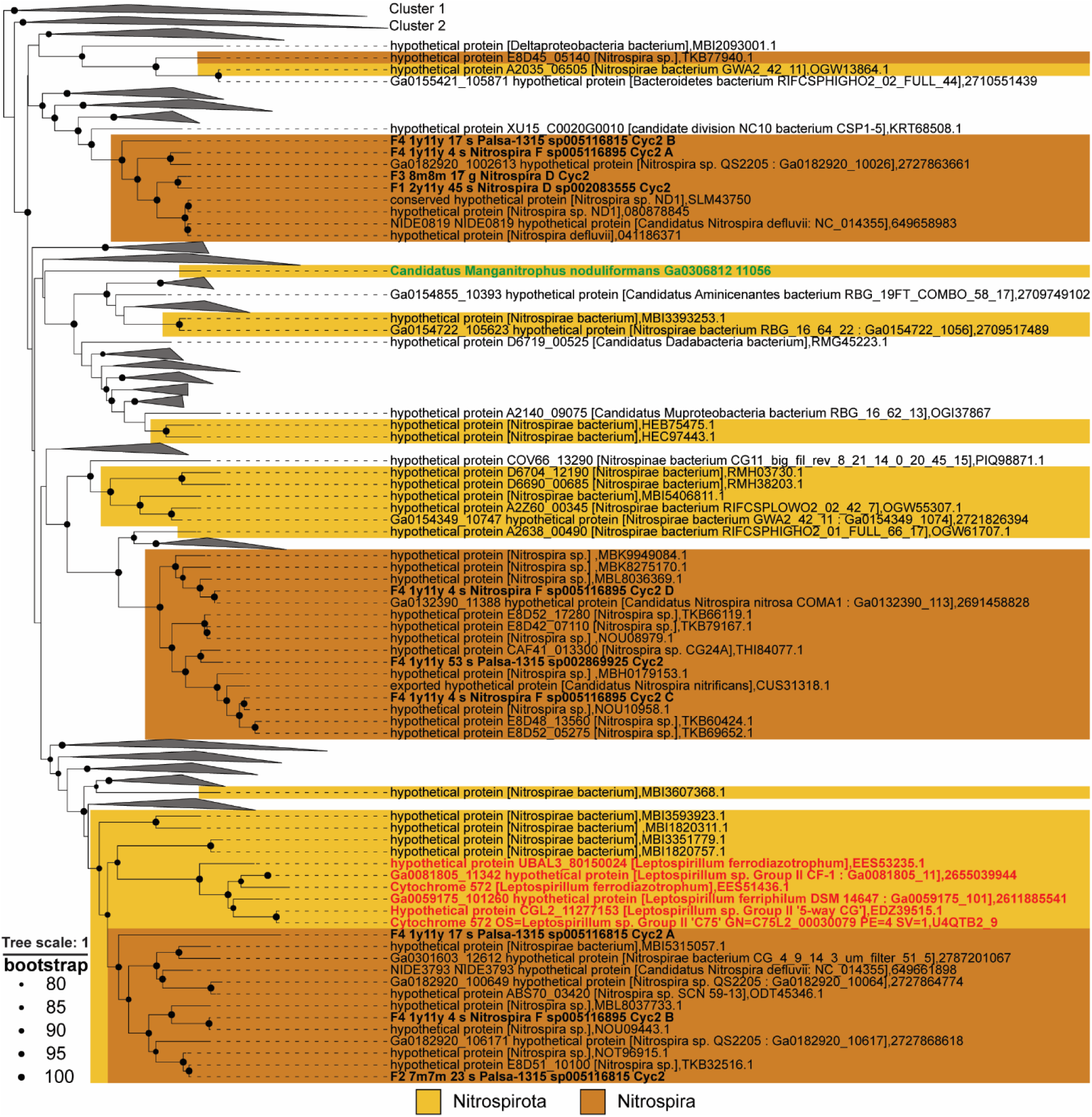
Phylogeny of Cyc2 proteins. The maximum-likelihood tree of 1626 Cyc2 sequences was rooted with cluster 1. Cluster 1, cluster 2, and all clades not containing *Nitrospirota* sequences within cluster 3 were collapsed. Sequences in MAGs identified in this study are highlighted in bold. Sequences with confirmed manganese- and iron-oxidizing functions are highlighted in green and red, respectively. A vector image of the full tree can be found in the supplementary materials (https://doi.org/10.5281/zenodo.14554862)

## 4. CONCLUSIONS

This study provides a comprehensive metagenomic analysis of the microbial communities within rapid sand filters, elucidating the roles of key taxa in biogeochemical processes associated with contaminant removal. A total of 34 high and 177 medium-quality MAGs were reconstructed, revealing functional insights into methane and iron oxidation, nitrification, and potential manganese oxidation.

Methanotrophic activity was primarily attributed to *Methylomonadaceae*, with dynamic shifts in dominant methanotrophic MAGs observed as filters aged. In younger filters, a MAG encoding the soluble methane monooxygenase alongside the particulate methane monooxygenase variants PMO and PXM dominated the methanotrophic community. However, in older filters, MAGs possessing only *pmoA* and *pxmA* genes became predominant, suggesting functional and ecological shifts over time, possibly caused by copper availability. Biological iron oxidation was restricted to the anthracite layer and linked to *Gallionella* MAGs that encoded cluster 1 Cyc2. *Gallionella* MAGs also became more abundant with filter age, indicating increasing relevance of biological iron oxidation in mature filters. Three MAGs affiliated with taxa never linked to iron oxidation (PHCI01*, Rubrivivax*, and *Candidatus* Nitrotoga) also contained cluster 1 *cyc2* genes, warranting further investigation into their iron-oxidizing capabilities.

The nitrifier community was dominated by AOB and canonical *Nitrospira* rather than comammox *Nitrospira*, challenging assumptions about comammox prevalence in rapid sand filters. Comammox *Nitrospira*, while present, only outnumbered canonical nitrifiers in the oldest sand layers, consistent with previous findings suggesting that they only become abundant after extended operational periods. The presence of manganese oxidation genes, such as *mnxG* and *moxA*, in *Nitrospira* MAGs, together with the apparent overrepresentation of canonical *Nitrospira* and the presence of cluster 3 Cyc2 homologs, suggests a role for *Nitrospira* in iron or manganese oxidation in these filters. Verification of this role will require targeted isolation and functional studies.

Overall, this metagenomic study advances our understanding of microbial community dynamics and their functional contributions in rapid sand filters treating groundwater for drinking water production. It highlights the intricate interplay of filter age, medium composition, and microbial metabolic potential in contaminant removal, offering new insights into previously unrecognized links between specific bacteria and the removal of certain contaminants, providing new targets for physiological studies to identify novel microbial players in contaminant removal.

## Supporting information

Supplementary_MAG_information

## 5. AUTHOR CONTRIBUTIONS

Conceptualization: AB, SH, PvdW, MvK, CS, SL

Supervision: PvdW, MvK, CS, SL

Sampling: AB, SH, FN

Investigation: AB, FN

Writing - Original Draft: AB

Writing - Review & Editing: PvdW, MvK, SL, CS, SH

Project administration: PvdW, MvK, CS, SL

Funding acquisition: SL, CS, MvK

## 6. DECLARATION OF COMPETING INTERESTS

The authors declare no competing interests.

## 7. DATA AVAILIBILITY

Supplementary data and figures have been submitted to the Zenodo repository under https://doi.org/10.5281/zenodo.14554862. Raw sequencing data and MAGs have been deposited in the NCBI Sequencing Read Archive (SRA) under BioProject PRJNA1200573

## 8. ACNKOWLEDGEMENTS

We are grateful to H. Doeve, M. Pipping, and Vitens N.V. for their support and collaboration during the visits to the drinking water treatment plants. We would like to thank Hanna Koch for her help with the NxrA/NarG alignement. This research was funded by the Netherlands Organisation for Scientific Research (NWO) partnership program Dunea–Vitens: Sand Filtration (grant 17841). MvK and SL were funded by NWO (016.Veni.192.062 and 016.Vidi.189.050, respectively), CP by the European Research Council (ERC Synergy Grant 694407 MARIX).

## Notes

https://doi.org/10.5281/zenodo.14554862

### Competing Interest Statement

The authors have declared no competing interest.

