## Supplementary tables and figures for "Metagenomic analysis of age-dependent microbial dynamics in dual-media rapid sand filters treating groundwater"

**SUPPLEMENTARY INFORMATION**

This PDF file contains:

Supplementary Figures S1-S3

Supplementary Table S1

Table S1. General information of all 34 high-quality and 177 medium-quality dereplicated MAGs.

| **MAG IDa** | **MAG taxonomyb** | **Length (Mb)** | **N50** | **# contigs** | **Compc** | **Contc** | **MIMAG qualityd** |
| --- | --- | --- | --- | --- | --- | --- | --- |
| F1_2y11y_1 | o_Bin18 | 6.41 | 22,100 | 498 | 95.34 | 7.52 | medium |
| F1_2y11y_2 | g_2-12-FULL-65-11 | 3.55 | 26,326 | 989 | 96.63 | 1.07 | medium |
| F1_2y11y_3 | g_OLB17 | 3.53 | 24,759 | 488 | 91.95 | 0.82 | medium |
| F1_2y11y_4 | g_JACMLD01 | 5.75 | 122,852 | 412 | 100 | 0 | medium |
| F1_2y11y_5 | g_WHSN01 | 3.10 | 6,465 | 360 | 86.58 | 3.4 | medium |
| F1_2y11y_7 | f_Planctomycetaceae | 4.39 | 5,022 | 1126 | 81.73 | 2.85 | medium |
| F1_2y11y_8 | s_JABFRY01_sp013140955 | 3.88 | 8,820 | 1024 | 99.52 | 0.53 | medium |
| F1_2y11y_9 | g_Hyphomicrobium | 4.37 | 14,052 | 1098 | 81.84 | 8.54 | medium |
| F1_2y11y_10 | g_DRPG01 | 10.10 | 13,205 | 903 | 84.35 | 4.81 | medium |
| F1_2y11y_11 | s_Nitrospira_A_sp900170025 | 4.05 | 12,314 | 897 | 99.81 | 1.92 | high |
| F1_2y11y_12 | g_RH-chloro-G1 | 2.71 | 10,153 | 66 | 84.83 | 2.62 | medium |
| F1_2y11y_13 | f_SXRF01 | 2.59 | 9,641 | 252 | 100 | 1.08 | medium |
| F1_2y11y_14 | g_CADEDH01 | 5.79 | 6,661 | 192 | 94.18 | 8.81 | medium |
| F1_2y11y_15 | g_Rubrivivax | 3.89 | 4,561 | 545 | 81.55 | 1.21 | medium |
| F1_2y11y_16 | f_Ahniellaceae | 3.69 | 4,136 | 235 | 85.55 | 1.32 | medium |
| F1_2y11y_17 | f_FEN-1173 | 2.17 | 2,695 | 594 | 73.87 | 2.58 | medium |
| F1_2y11y_18 | f_GCA-2862545 | 8.40 | 14,580 | 88 | 95.14 | 4.31 | medium |
| F1_2y11y_19 | g_Terricaulis | 4.04 | 141,248 | 1206 | 99.91 | 0.61 | high |
| F1_2y11y_20 | g_2-02-FULL-65-29 | 2.76 | 20,043 | 1220 | 97.96 | 2.52 | medium |
| F1_2y11y_21 | g_Gallionella | 2.18 | 5,116 | 86 | 96.03 | 8.1 | medium |
| F1_2y11y_22 | g_CADEEN01 | 3.74 | 34,074 | 210 | 98.34 | 0.1 | medium |
| F1_2y11y_23 | g_CAIUGU01 | 3.67 | 11,316 | 62 | 86.52 | 1.51 | medium |
| F1_2y11y_24 | g_SCMT01 | 4.38 | 135,448 | 233 | 99.99 | 0.01 | medium |
| F1_2y11y_25 | g_WHSN01 | 4.47 | 4,734 | 302 | 100 | 4.56 | medium |
| F1_2y11y_26 | g_JADJLQ01 | 5.31 | 6,421 | 1552 | 97.17 | 4.05 | medium |
| F1_2y11y_27 | g_Nitrosomonas | 3.22 | 85,111 | 142 | 99.98 | 0.02 | medium |
| F1_2y11y_28 | g_JACRMR01 | 3.20 | 23,836 | 859 | 95.05 | 0.04 | high |
| F1_2y11y_29 | g_SCMT01 | 4.17 | 129,843 | 417 | 99.99 | 0.14 | medium |
| F1_2y11y_30 | g_2-02-FULL-65-29 | 5.98 | 32,649 | 222 | 100 | 0.23 | medium |
| F1_2y11y_31 | o_Fen-336 | 7.42 | 6,122 | 2792 | 95.13 | 4.77 | high |
| F1_2y11y_32 | s_UBA7657_sp002483465 | 2.04 | 18,887 | 2508 | 97.55 | 3.05 | high |
| F1_2y11y_33 | s_Gp6-AA40_sp013140775 | 5.02 | 8,124 | 1145 | 90.61 | 7.91 | medium |
| F1_2y11y_34 | c_Mor1 | 4.06 | 14,939 | 741 | 95.74 | 1.03 | high |
| F1_2y11y_35 | g_WHSN01 | 3.82 | 32,035 | 102 | 99.58 | 1.02 | medium |
| F1_2y11y_36 | g_JAEUJP01 | 10.25 | 4,178 | 453 | 92.76 | 6.09 | medium |
| F1_2y11y_37 | g_DRPG01 | 9.28 | 4,328 | 242 | 89.21 | 9.33 | medium |
| F1_2y11y_38 | f_JAGNNV01 | 4.49 | 4,629 | 794 | 84.04 | 2.73 | medium |
| F1_2y11y_39 | f_UBA955 | 2.61 | 3,812 | 619 | 79.67 | 5.88 | medium |
| F1_2y11y_40 | g_SPCO01 | 3.02 | 8,982 | 127 | 85.79 | 3.34 | medium |
| F1_2y11y_41 | g_Nitrosomonas | 2.47 | 17,507 | 451 | 99.99 | 2.02 | medium |
| F1_2y11y_42 | g_DSQF01 | 3.52 | 5,579 | 451 | 91.19 | 4.35 | medium |
| F1_2y11y_43 | g_Sphingorhabdus_B | 2.70 | 5,308 | 568 | 84.53 | 5.56 | medium |
| F1_2y11y_44 | g_Novosphingobium | 2.93 | 34,195 | 1125 | 99.99 | 0 | medium |
| F1_2y11y_45 | s_Nitrospira_D_sp002083555 | 3.21 | 10,883 | 587 | 86.93 | 2.75 | medium |
| F1_2y11y_46 | c_RBG-16-71-46 | 3.60 | 10,952 | 456 | 94.91 | 0.31 | medium |
| F2_7m7m_1 | o_Planctomycetales | 5.92 | 23,301 | 396 | 96.69 | 0.47 | medium |
| F2_7m7m_2 | g_Flavobacterium | 2.67 | 12,347 | 23 | 93.31 | 0.14 | medium |
| F2_7m7m_3 | g_PHCI01 | 4.59 | 6,518 | 342 | 95.31 | 6.75 | medium |
| F2_7m7m_4 | g_Sphingopyxis | 2.16 | 3,579 | 493 | 85.15 | 3.13 | medium |
| F2_7m7m_5 | g_GWA2-66-18 | 2.66 | 3,085 | 635 | 82.03 | 2.5 | medium |
| F2_7m7m_6 | g_Pseudolysinimonas | 2.15 | 2,800 | 559 | 81.33 | 3.31 | medium |
| F2_7m7m_7 | s_UBA7672_sp016714495 | 4.22 | 20,193 | 564 | 99.07 | 3.26 | medium |
| F2_7m7m_8 | g_ELB16-189 | 5.27 | 15,458 | 79 | 92.62 | 1.61 | medium |
| F2_7m7m_9 | f_Zambryskibacteraceae | 1.44 | 48,435 | 285 | 100 | 2.58 | high |
| F2_7m7m_10 | g_SCMT01 | 4.25 | 298,187 | 731 | 100 | 0.93 | medium |
| F2_7m7m_11 | g_SHVO01 | 3.69 | 15,468 | 225 | 96.71 | 3.74 | medium |
| F2_7m7m_12 | g_RBG-16-66-20 | 3.14 | 8,465 | 308 | 90.63 | 2.21 | medium |
| F2_7m7m_13 | g_Arenimonas | 3.07 | 6,224 | 1106 | 87.27 | 3.13 | medium |
| F2_7m7m_14 | g_CAIKVZ01 | 2.73 | 6,756 | 398 | 97.84 | 2.53 | medium |
| F2_7m7m_15 | f_Chthoniobacteraceae | 5.14 | 11,002 | 1096 | 88.19 | 0.85 | medium |
| F2_7m7m_16 | g_ELB16-189 | 5.08 | 154,932 | 579 | 99.5 | 0.19 | medium |
| F2_7m7m_17 | g_SZUA-430 | 4.54 | 23,633 | 1779 | 98.02 | 0.26 | medium |
| F2_7m7m_18 | g_Gallionella | 2.13 | 3,524 | 135 | 83.1 | 2.9 | medium |
| F2_7m7m_19 | s_JADJWH01_sp016712635 | 2.97 | 23,224 | 144 | 97.65 | 0.29 | medium |
| F2_7m7m_20 | g_VFZH01 | 3.34 | 3,738 | 822 | 96.22 | 2.71 | medium |
| F2_7m7m_21 | g_UBA5216 | 4.21 | 16,718 | 27 | 92.37 | 4.06 | medium |
| F2_7m7m_22 | g_DRPG01 | 8.57 | 10,365 | 1002 | 93.76 | 6.98 | medium |
| F2_7m7m_23 | s_Palsa-1315_sp005116815 | 1.94 | 4,363 | 121 | 71.57 | 2.95 | medium |
| F2_7m7m_24 | s_JABFRJ01_sp013141255 | 4.18 | 2,557 | 236 | 78.57 | 8.17 | medium |
| F2_7m7m_26 | g_Rubrivivax | 5.93 | 118,938 | 595 | 99.92 | 1.05 | medium |
| F2_7m7m_27 | g_Devosia_A | 3.44 | 47,547 | 765 | 93.49 | 0.39 | medium |
| F2_7m7m_28 | g_UBA5216 | 4.32 | 8,471 | 1407 | 96.42 | 7.79 | medium |
| F2_7m7m_29 | o_Bdellovibrionales | 4.04 | 462,047 | 332 | 100 | 2.73 | high |
| F2_7m7m_30 | g_Flavobacterium | 3.96 | 79,984 | 564 | 100 | 0.35 | high |
| F2_7m7m_31 | g_Aestuariivirga | 3.61 | 31,647 | 1000 | 95.17 | 1.44 | medium |
| F2_7m7m_32 | g_SHZI01 | 3.10 | 6,822 | 870 | 92.4 | 4.9 | medium |
| F2_7m7m_33 | g_Methyloglobulus | 3.18 | 5,042 | 8 | 94.75 | 4.75 | medium |
| F2_7m7m_34 | f_SIAN01 | 4.32 | 3,314 | 718 | 78.72 | 4.75 | medium |
| F2_7m7m_35 | g_LMDS01 | 5.94 | 26,793 | 52 | 99.18 | 1.28 | medium |
| F2_7m7m_36 | g_JAEUMZ01 | 3.40 | 7,971 | 28 | 95.2 | 2.56 | medium |
| F2_7m7m_37 | f_Kaistiaceae | 3.35 | 3,839 | 911 | 79.25 | 9.86 | medium |
| F2_7m7m_38 | f_UBA2999 | 4.31 | 6,333 | 662 | 92.31 | 6.42 | medium |
| F2_7m7m_39 | o_UBA9983_A | 0.71 | 123,190 | 878 | 96.32 | 0.01 | high |
| F2_7m7m_40 | g_REEB506 | 2.92 | 102,089 | 92 | 99.92 | 0.2 | medium |
| F2_7m7m_41 | f_Zambryskibacteraceae | 0.74 | 77,093 | 77 | 100 | 1.21 | high |
| F2_7m7m_42 | g_M3007 | 5.55 | 7,984 | 205 | 98.57 | 0.54 | medium |
| F2_7m7m_43 | g_Aestuariivirga | 3.05 | 5,849 | 812 | 80.58 | 0.5 | medium |
| F2_7m7m_44 | g_CADEFD01 | 5.89 | 9,532 | 787 | 90.44 | 1.01 | medium |
| F2_7m7m_45 | g_VFBF01 | 4.33 | 73,713 | 995 | 100 | 0.16 | medium |
| F2_7m7m_46 | g_OLB19 | 0.81 | 27,497 | 7 | 98.21 | 3.1 | high |
| F2_7m7m_47 | g_WHSN01 | 5.73 | 58,252 | 857 | 100 | 2.55 | medium |
| F2_7m7m_48 | g_Sediminibacterium | 3.22 | 4,596 | 320 | 92.29 | 0.06 | medium |
| F2_7m7m_49 | g_JAEUPI01 | 4.92 | 8,239 | 440 | 96.73 | 2 | high |
| F2_7m7m_50 | g_C7867-006 | 0.83 | 214,889 | 93 | 100 | 0.45 | high |
| F3_2m2m_1 | g_Methylotenera | 2.22 | 8,646 | 341 | 77.79 | 7.56 | medium |
| F3_2m2m_2 | g_UBA1573 | 2.12 | 24,757 | 108 | 98.76 | 2.33 | high |
| F3_2m2m_3 | g_JAAFJR01 | 3.91 | 34,419 | 56 | 97.67 | 6.06 | medium |
| F3_2m2m_4 | g_JABFSH01 | 3.81 | 18,258 | 19 | 85.75 | 0.11 | medium |
| F3_2m2m_5 | g_Methylotenera | 2.69 | 55,014 | 31 | 96.25 | 2.51 | medium |
| F3_2m2m_6 | s_Methylotenera_sp013141495 | 2.18 | 15,175 | 244 | 89.03 | 1.44 | medium |
| F3_2m2m_7 | f_Methylophilaceae | 2.72 | 37,091 | 154 | 96.31 | 2.69 | medium |
| F3_2m2m_8 | g_Albitalea | 3.69 | 7,916 | 317 | 94.22 | 4.5 | medium |
| F3_2m2m_9 | g_Gallionella | 2.23 | 4,102 | 67 | 85.12 | 2.35 | medium |
| F3_2m2m_11 | o_RUG11792 | 1.38 | 24,783 | 554 | 100 | 0.85 | high |
| F3_2m2m_12 | g_SCTK01 | 2.69 | 104,626 | 152 | 100 | 0.3 | medium |
| F3_2m2m_13 | f_SXRF01 | 2.06 | 141,876 | 80 | 100 | 0 | high |
| F3_2m2m_14 | g_Rhabdaerophilum | 3.39 | 218,784 | 252 | 98.87 | 0.15 | high |
| F3_2m2m_15 | s_Nitrosomonas_sp902826115 | 3.83 | 42,242 | 1442 | 100 | 5.7 | medium |
| F3_2m2m_16 | g_Nitrosomonas | 2.34 | 23,587 | 33 | 93.99 | 0 | medium |
| F3_2m2m_17 | f_SXRF01 | 2.12 | 9,508 | 512 | 95.68 | 1.34 | medium |
| F3_2m2m_18 | g_UBA5078 | 3.42 | 149,240 | 349 | 99.77 | 0.58 | medium |
| F3_2m2m_19 | g_Rubrivivax | 5.91 | 19,103 | 226 | 97.69 | 3 | medium |
| F3_2m2m_20 | f_UBA3002 | 2.22 | 49,805 | 266 | 99.4 | 0.04 | high |
| F3_2m2m_21 | g_UBA1942 | 2.34 | 11,386 | 273 | 86.98 | 0.58 | medium |
| F3_2m2m_22 | s_Escherichia_coli | 3.34 | 2,612 | 1022 | 70.71 | 2.26 | medium |
| F3_2m2m_23 | f_CG1-02-43-26 | 1.74 | 115,458 | 270 | 95.57 | 0 | high |
| F3_2m2m_24 | g_LMDS01 | 2.69 | 6,211 | 262 | 82.74 | 2 | medium |
| F3_2m2m_25 | g_SXIZ01 | 4.60 | 18,159 | 111 | 92.92 | 2.18 | medium |
| F3_2m2m_26 | g_Tabrizicola | 3.98 | 29,782 | 125 | 100 | 0.14 | medium |
| F3_2m2m_27 | g_Fluviicola | 3.33 | 15,885 | 466 | 88.23 | 0.35 | medium |
| F3_2m2m_28 | s_Methylotenera_sp013141075 | 2.36 | 15,210 | 269 | 95.49 | 0.38 | medium |
| F3_2m2m_29 | g_Legionella | 3.27 | 4,502 | 691 | 91.94 | 8.74 | medium |
| F3_2m2m_30 | g_Nitrosomonas | 3.10 | 21,589 | 74 | 97.64 | 5.96 | medium |
| F3_2m2m_31 | f_UBA11393 | 1.82 | 56,735 | 161 | 100 | 4.13 | medium |
| F3_2m2m_32 | f_Micavibrionaceae | 2.06 | 25,587 | 371 | 99.56 | 0.49 | medium |
| F3_2m2m_34 | g_Asticcacaulis | 2.34 | 6,576 | 318 | 94.76 | 1.64 | medium |
| F3_2m2m_35 | g_Ga0077554 | 3.04 | 24,028 | 332 | 98.68 | 1.09 | medium |
| F3_2m2m_36 | s_Methylovulum_psychrotolerans | 4.38 | 8,747 | 101 | 87.93 | 4.16 | medium |
| F3_2m2m_37 | g_UBA1942 | 2.82 | 69,314 | 277 | 99.4 | 0.33 | medium |
| F3_2m2m_38 | g_Hylemonella | 3.79 | 56,646 | 127 | 99.79 | 0.48 | medium |
| F3_2m2m_39 | g_Rubrivivax | 4.78 | 21,294 | 499 | 98.96 | 0.25 | medium |
| F3_2m2m_40 | s_Methylotenera_mobilis_B | 2.66 | 13,888 | 117 | 97.67 | 4.51 | medium |
| F3_2m2m_41 | g_Hylemonella | 3.86 | 199,696 | 211 | 99.14 | 4.88 | medium |
| F3_2m2m_42 | f_SXRF01 | 2.32 | 58,568 | 95 | 100 | 3.65 | high |
| F3_2m2m_43 | g_SXIZ01 | 3.60 | 91,507 | 639 | 99.99 | 0.19 | high |
| F3_2m2m_44 | f_Nocardioidaceae | 2.47 | 6,340 | 668 | 92.46 | 1.65 | medium |
| F3_8m8m_1 | g_C7867-006 | 0.66 | 17,337 | 61 | 92.64 | 0.93 | medium |
| F3_8m8m_2 | g_Nitrosomonas | 2.50 | 44,170 | 561 | 99.94 | 0 | high |
| F3_8m8m_3 | g_M3007 | 5.57 | 8,405 | 368 | 89.08 | 1.43 | medium |
| F3_8m8m_5 | g_CAIXRL01 | 4.67 | 6,428 | 1136 | 93.47 | 0.86 | medium |
| F3_8m8m_6 | g_UBA4660 | 4.08 | 9,033 | 969 | 92.51 | 0.16 | medium |
| F3_8m8m_7 | g_Arenimonas | 3.53 | 24,009 | 802 | 99.99 | 1.15 | medium |
| F3_8m8m_8 | g_Gallionella | 1.67 | 3,044 | 302 | 82.11 | 5.61 | medium |
| F3_8m8m_9 | g_Methylotenera_A | 2.45 | 139,841 | 33 | 99.74 | 0.04 | medium |
| F3_8m8m_10 | g_Gallionella | 2.26 | 5,883 | 72 | 93.22 | 8.4 | medium |
| F3_8m8m_11 | g_Nitrotoga | 2.83 | 13,004 | 215 | 100 | 2.45 | medium |
| F3_8m8m_12 | g_SCTL01 | 4.35 | 5,465 | 316 | 90.77 | 8.87 | medium |
| F3_8m8m_13 | g_LMDS01 | 3.71 | 4,385 | 106 | 78.89 | 3.78 | medium |
| F3_8m8m_14 | c_PLA2 | 4.58 | 7,361 | 6 | 85.55 | 1.48 | medium |
| F3_8m8m_15 | g_Novosphingobium | 4.45 | 24,791 | 640 | 96.92 | 1.01 | medium |
| F3_8m8m_16 | f_UBA9973 | 0.65 | 34,066 | 790 | 100 | 0.05 | high |
| F3_8m8m_17 | g_Nitrospira_D | 3.67 | 93,490 | 813 | 98.97 | 0.33 | medium |
| F3_8m8m_18 | g_UBA7672 | 4.20 | 30,978 | 1060 | 92.26 | 0.74 | medium |
| F3_8m8m_19 | g_JADKGY01 | 5.27 | 32,342 | 561 | 97.65 | 0.38 | medium |
| F3_8m8m_20 | g_UBA910 | 0.75 | 274,308 | 241 | 99.66 | 0.36 | high |
| F3_8m8m_21 | g_SHZI01 | 3.35 | 7,352 | 663 | 93.26 | 5.95 | medium |
| F3_8m8m_22 | g_SZUA-430 | 4.09 | 6,676 | 77 | 90.25 | 1.51 | medium |
| F4_1y11y_1 | g_Terricaulis | 2.58 | 7,515 | 415 | 75.67 | 1.83 | medium |
| F4_1y11y_2 | f_SG-bin7 | 3.67 | 47,803 | 1416 | 93.65 | 0.17 | medium |
| F4_1y11y_3 | g_Gp6-AA40 | 3.40 | 45,195 | 989 | 97.39 | 0.57 | medium |
| F4_1y11y_4 | s_Nitrospira_F_sp005116895 | 3.97 | 56,714 | 47 | 98.89 | 0.77 | medium |
| F4_1y11y_5 | f_Kaistiaceae | 3.05 | 45,788 | 1582 | 97.15 | 0.08 | medium |
| F4_1y11y_6 | g_Terricaulis | 3.34 | 57,231 | 86 | 99.86 | 0.57 | high |
| F4_1y11y_7 | s_JABFRJ01_sp013141255 | 4.54 | 13,330 | 114 | 94.99 | 1.56 | medium |
| F4_1y11y_8 | g_LMDS01 | 5.70 | 248,437 | 222 | 99.04 | 3.96 | medium |
| F4_1y11y_9 | g_JAAFIP01 | 3.64 | 180,702 | 488 | 99.93 | 0.42 | high |
| F4_1y11y_10 | f_UBA955 | 3.37 | 2,672 | 137 | 74.14 | 3.8 | medium |
| F4_1y11y_11 | g_UBA668 | 3.10 | 3,831 | 440 | 93.59 | 2.71 | medium |
| F4_1y11y_12 | g_Sphingopyxis | 2.35 | 113,674 | 133 | 99.86 | 0.42 | high |
| F4_1y11y_13 | f_JAGNNV01 | 4.67 | 3,474 | 1588 | 90.18 | 3.93 | medium |
| F4_1y11y_14 | f_UBA2999 | 4.20 | 188,801 | 1015 | 100 | 0.21 | high |
| F4_1y11y_15 | f_JAFNAJ01 | 5.94 | 171,070 | 2592 | 100 | 0.68 | high |
| F4_1y11y_16 | g_RBC038 | 2.97 | 24,338 | 45 | 94.09 | 1.39 | medium |
| F4_1y11y_17 | s_Palsa-1315_sp005116815 | 2.00 | 5,472 | 327 | 85.48 | 4.18 | medium |
| F4_1y11y_18 | p_ARS69 | 4.10 | 76,903 | 63 | 95.26 | 2.11 | high |
| F4_1y11y_19 | g_JAEUIV01 | 3.50 | 15,437 | 300 | 91.55 | 2.55 | medium |
| F4_1y11y_20 | f_JADIYM01 | 5.60 | 4,554 | 882 | 91.58 | 3.25 | medium |
| F4_1y11y_21 | g_UBA2982 | 3.07 | 3,673 | 416 | 89.13 | 0.54 | medium |
| F4_1y11y_23 | g_SXOA01 | 6.37 | 2,878 | 156 | 77.43 | 4.19 | medium |
| F4_1y11y_24 | g_SCMT01 | 4.62 | 521,729 | 56 | 100 | 0.68 | medium |
| F4_1y11y_25 | g_Sphingorhabdus_B | 3.20 | 27,621 | 171 | 97.34 | 4.77 | medium |
| F4_1y11y_26 | g_Arenimonas | 3.48 | 386,094 | 544 | 99.98 | 0.06 | medium |
| F4_1y11y_27 | g_CAIUGU01 | 4.35 | 28,228 | 1164 | 95.88 | 4.54 | high |
| F4_1y11y_28 | s_Methylotenera_sp013141075 | 1.91 | 2,400 | 866 | 83.05 | 4.92 | medium |
| F4_1y11y_29 | f_JAGNNV01 | 3.77 | 13,546 | 668 | 95.92 | 3.34 | medium |
| F4_1y11y_30 | s_Novosphingobium_sp013141325 | 2.56 | 71,081 | 386 | 96.39 | 0.05 | medium |
| F4_1y11y_31 | f_Hyphomicrobiaceae | 4.79 | 52,744 | 871 | 99.45 | 1.03 | medium |
| F4_1y11y_32 | g_CADEFD01 | 6.03 | 20,444 | 938 | 100 | 3.68 | high |
| F4_1y11y_33 | f_Pseudonocardiaceae | 5.70 | 5,999 | 545 | 92.19 | 2.84 | medium |
| F4_1y11y_34 | f_WHTK01 | 3.89 | 5,509 | 140 | 86.83 | 5.83 | medium |
| F4_1y11y_35 | g_Novosphingobium | 4.26 | 8,911 | 711 | 98.92 | 1.8 | medium |
| F4_1y11y_36 | f_JAGNNV01 | 4.42 | 18,798 | 850 | 98.03 | 3.6 | medium |
| F4_1y11y_37 | g_Anatilimnocola | 7.30 | 12,611 | 833 | 98.17 | 4.73 | high |
| F4_1y11y_38 | f_JAEUIK01 | 5.59 | 8,305 | 282 | 93.78 | 0.21 | medium |
| F4_1y11y_39 | g_Terricaulis | 3.47 | 8,573 | 244 | 86.57 | 9.09 | medium |
| F4_1y11y_40 | f_Tepidiformaceae | 4.48 | 8,885 | 842 | 87.95 | 5.29 | medium |
| F4_1y11y_41 | g_Nitrotoga | 2.69 | 3,711 | 477 | 87.74 | 9.26 | medium |
| F4_1y11y_42 | g_Aestuariivirga | 3.81 | 5,861 | 274 | 89 | 4.45 | medium |
| F4_1y11y_43 | g_Gallionella | 2.39 | 15,117 | 664 | 88.21 | 3.46 | medium |
| F4_1y11y_44 | f_Kaistiaceae | 3.64 | 31,347 | 235 | 93.94 | 3.21 | medium |
| F4_1y11y_45 | g_JAEUPI01 | 5.04 | 8,272 | 152 | 88.92 | 4.22 | medium |
| F4_1y11y_46 | g_Methyloglobulus | 3.60 | 11,325 | 1032 | 94.19 | 0.92 | medium |
| F4_1y11y_47 | g_Hyphomicrobium | 4.30 | 27,723 | 451 | 95.24 | 0.68 | medium |
| F4_1y11y_48 | g_Methyloglobulus | 4.09 | 8,168 | 96 | 96.54 | 8.28 | medium |
| F4_1y11y_49 | g_WHSN01 | 4.37 | 36,065 | 111 | 100 | 1.9 | medium |
| F4_1y11y_50 | g_WHSN01 | 6.00 | 7,433 | 352 | 99.61 | 7.7 | medium |
| F4_1y11y_51 | g_SCUD01 | 2.91 | 9,556 | 488 | 93.15 | 4.72 | medium |
| F4_1y11y_52 | g_SPCO01 | 3.23 | 67,360 | 94 | 93.06 | 0.6 | medium |
| F4_1y11y_53 | s_Palsa-1315_sp002869925 | 4.33 | 75,378 | 482 | 99.99 | 0.34 | medium |
| F4_1y11y_54 | o_UBA2979 | 2.50 | 11,121 | 69 | 90.09 | 1.72 | medium |
| F4_1y11y_55 | f_Hyphomicrobiaceae | 4.15 | 12,411 | 33 | 94.87 | 3.45 | medium |

^a^ MAG ID is comprised of the assembly of origin followed by an arbitrary number.
^b^ Taxonomy as determined by GTDKtk v2.1.1 (Chaumeil et al. 2022), where the smaller case letter represents the furthest determined taxonomic level up until genus.
^c^ Completeness (Comp) and contamination (Cont) were calculated using CheckM2 v0.1.3 (Chklovski et al. 2023).
^d^ High and medium-quality MAGs were defined as having completeness >90 % and contamination <5 %, and completeness >70 % and contamination <10 %, respectively. Additionally, high-quality MAGs must contain 5S, 16S, and 23S genes and at least 18 tRNAs.

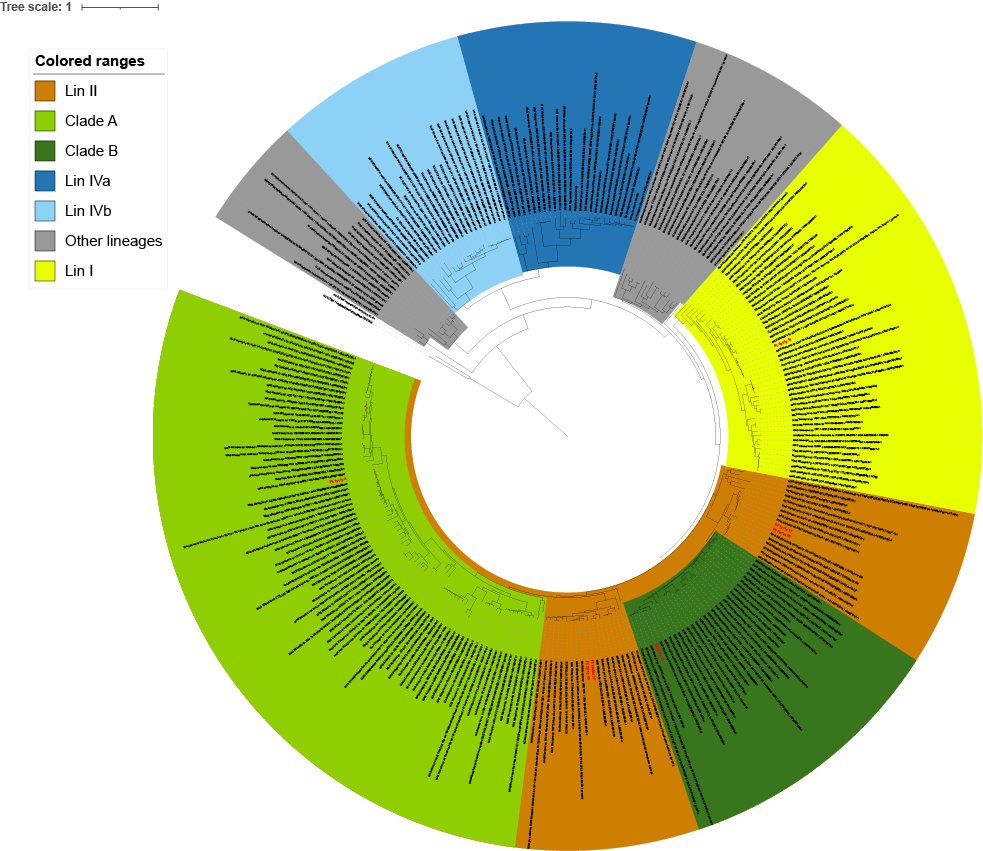

Figure S1. Whole genome phylogenetic tree of *Nitrospira*. An concatenated alignment of 92 single-core copy genes of the 7 *Nitrospira* MAGs identified in this study along with a reference database of 265 genomes was made using UBCG. A maximum-likelihood tree was then generated using IQ-TREE v2.1.4 using the GTR+F+I+G4 model and 1000 ultrafast bootstraps. The tree is rooted using two *Leptospirillum* genomes as outgroup. Different *Nitrospira* lineages and comammox clades are visualized using colored ranges. A vector image of the full tree can be found in the supplementary materials (https://doi.org/10.5281/zenodo.14554862)

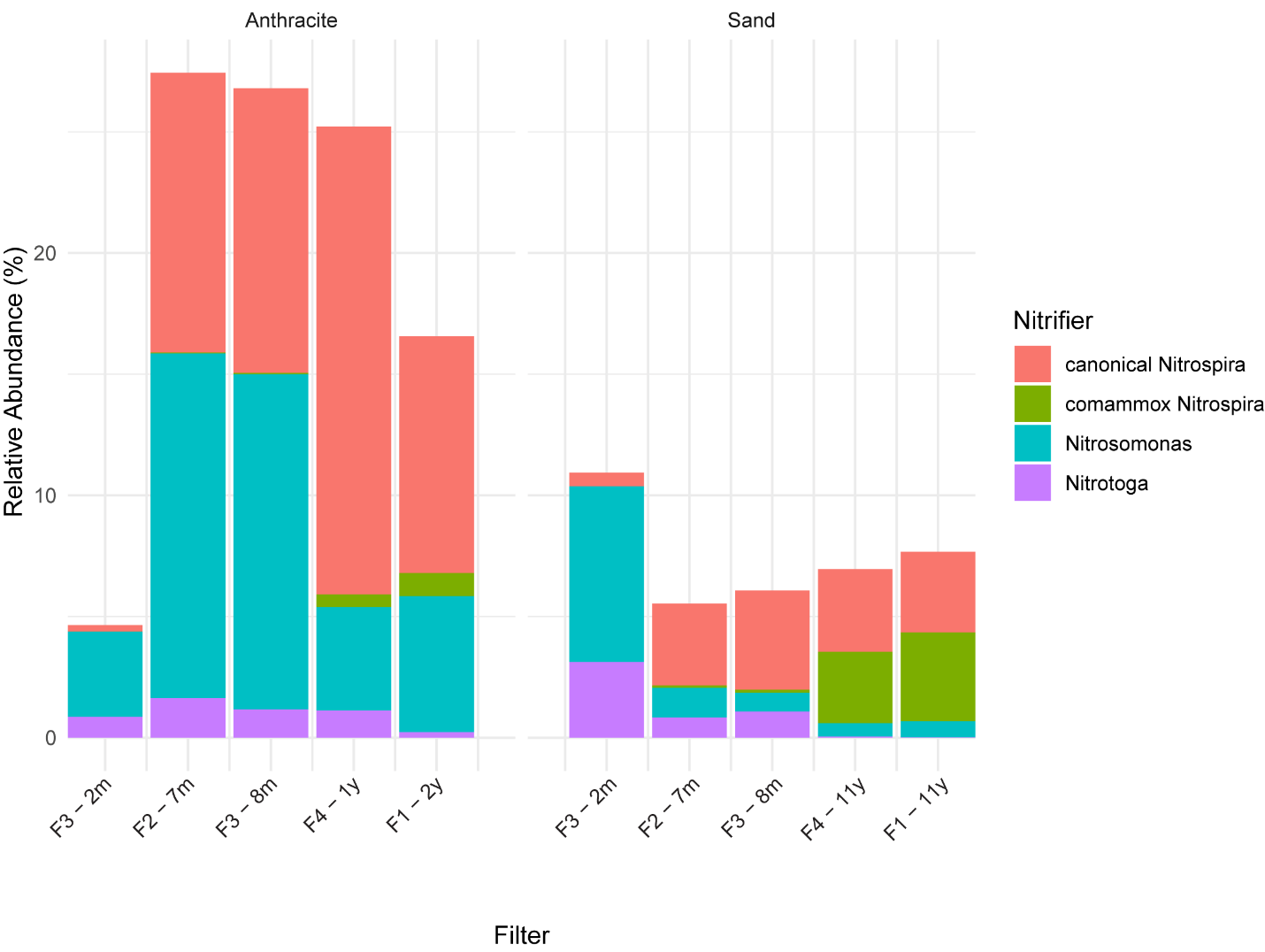

Figure S2. Relative abundance of nitrifier MAGs. m = months and y = years

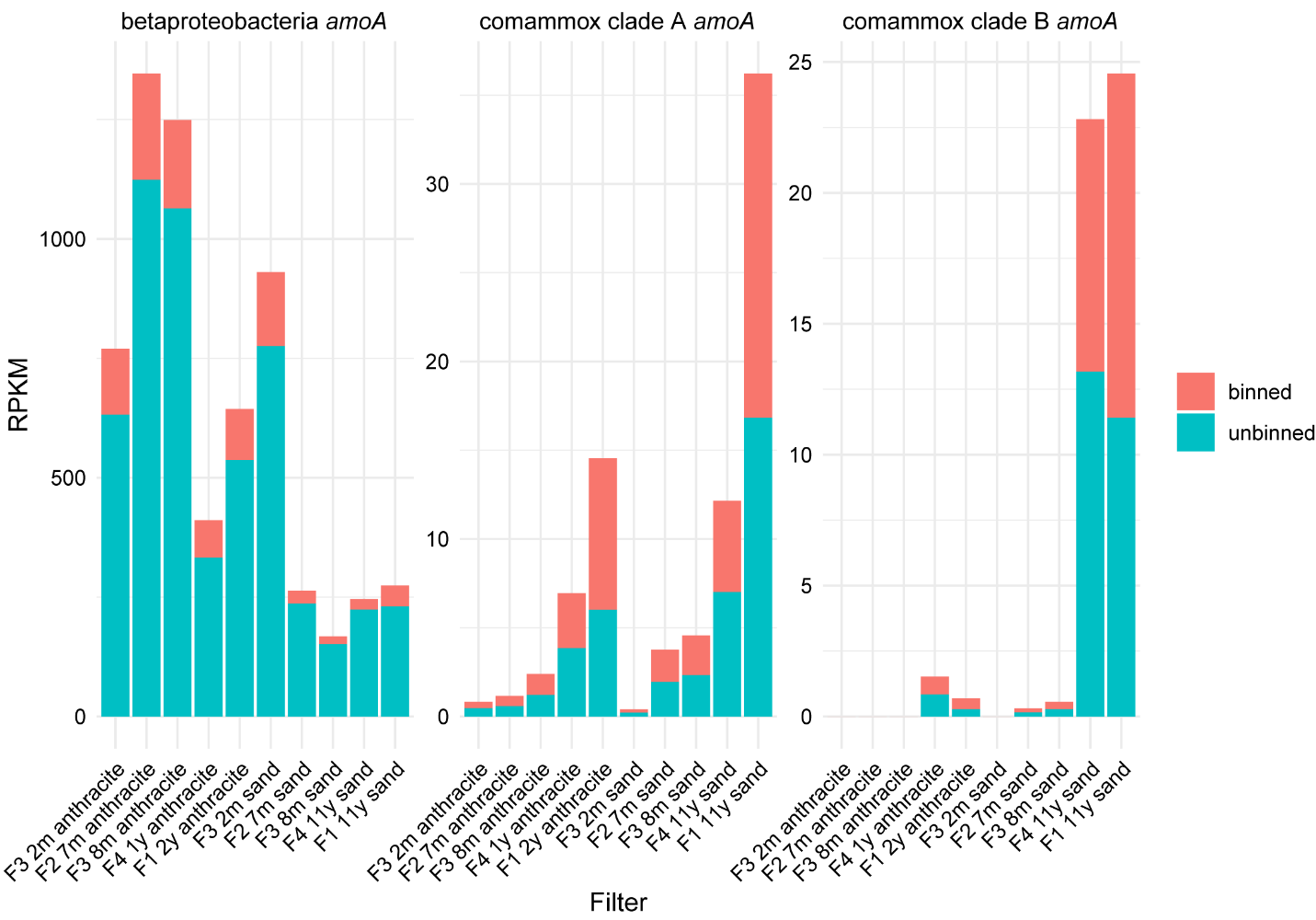

Figure S3. Abundance of binned and unbinned *amoA* genes.
